# Evolution and maintenance of microbe-mediated protection under occasional pathogen infection

**DOI:** 10.1101/2020.01.24.917138

**Authors:** A. Kloock, M.B. Bonsall, K.C. King

## Abstract

Every host is colonized by a variety of microbes, some of which can protect their hosts from pathogen infection. However, pathogen presence naturally varies over time in nature, such as in the case of seasonal epidemics. We experimentally coevolved populations of *Caenorhabditis elegans* worm hosts with bacteria possessing protective traits (*Enterococcus faecalis*), in treatments varying the infection frequency with pathogenic *Staphylococcus aureus* every host generation, alternating host generations, every fifth host generation or never. We additionally investigated the effect of initial pathogen presence at the formation of the defensive symbiosis. Our results show that enhanced microbe-mediated protection evolved during host-protective microbe coevolution when faced with rare infections by a pathogen. Initial pathogen presence had no effect on the evolutionary outcome of microbe-mediated protection. We also found that protection was only effective at preventing mortality during the time of pathogen infection. Overall, our results suggest that resident microbes can be a form of transgenerational immunity against rare pathogen infection.

## Introduction

In nature, all plants and animals are colonised by microbes (Barriere, 2006; Ley et al., 2006; Vántus et al., 2014). The composition of these microbial communities is highly diverse and includes harmful, neutral, and beneficial microbial species (Ley et al., 2006), including those that can be important players in host defence against parasites, a phenomenon referred to as ‘defensive mutualism’ (King, 2019; May and Nelson, 2014). Recognised for over a century, defensive mutualism has been observed in plants (Mendes et al., 2011) and in a range of animals (Dillon et al., 2000; Dong et al., 2009; Jaenike et al., 2010; Koch and Schmid-Hempel, 2011), including humans (Kamada et al., 2013; Ley et al., 2006; Maynard et al., 2012) wherein microbes can supplement host immune systems (Abt and Artis, 2013; Hooper et al., 2012; McFall-Ngai et al., 2013).

The net benefits of defensive mutualism are dependent upon the presence of pathogens (Clay et al., 2005; King and Bonsall, 2017; Lively et al., 2005). Whilst hosts can benefit from microbe-mediated protection, defensive symbionts can be less beneficial to the host in the absence of enemies, due to metabolic and physiological costs (King, 2019). For example, in the interaction of aphids and the bacterium *Hamiltonella defensa*, the host tissue is harmed by defensive toxins that protect against infection from parasitoids (Vorburger and Gouskov, 2011). In some cases, possessing protective microbes might be more beneficial to the host than investing in its own immune system (Martinez et al., 2016). From the perspective of the symbiont, it is most useful to its host under high pathogen prevalence, and thus can persist in the host population (Palmer et al., 2008). Nevertheless, a stable symbiotic interaction is hypothesized to be evolved and maintained (Kwiatkowski and Vorburger, 2012) only when the host benefit of carrying defensive symbionts outweighs any costs. The interactions of obligate and defensive symbionts and hosts can be stable for millions of years (Moran et al., 2005).

Not all environments are constantly pathogen rich which might shift the balance of costs and benefits during defensive mutualisms, particularly during coevolutionary interactions (King and Bonsall, 2017). Pathogen prevalence can be spatially (King et al., 2009) or temporally variable, the latter in the case of seasonal epidemics (e.g., flu peaks each winter in the northern hemisphere (Finkelman, 2007) or rabies in North American skunks which peaks in Autumn (Gremillion-Smith and Woolf, 1988)). Different environmental factors can influence disease transmission such as an increase in malaria risk in warmer regions after rainfall (Altizer et al., 2006), or an increase in contact rate and thus higher flu infection rate during the winter months (London and Yorke, 1973). The impact of other temporally heterogeneous factors on the strength and direction of selection on species interactions have been explored (oxygen concentration (Dey et al., 2016), resource availability (Friman et al., 2011; Friman and Laakso, 2011; Hiltunen et al., 2012), environmental productivity (Harrison et al., 2013)). Whether the varied presence of pathogens can similarly alter selection for symbiotic interactions has been explored theoretically (Fenton et al., 2011), but remains to be empirically tested.

Here, we examined the impact of temporal variation in pathogen infection on the evolution of microbe-mediated protection. We used *Caenorhabditis elegans* as a worm host and allowed it to be colonised by a bacterium (*Enterococcus faecalis*) that protects against infection by *Staphylococcus aureus* (King et al., 2016). *Enterococcus faecalis* has been shown to be protective across animal microbiomes (Kommineni et al., 2015; Martín-Vivaldi et al., 2010). It has been previously shown that *E. faecalis* can evolve to provide enhanced protection when residing in *C. elegans* hosts during constant pathogen infection (King et al., 2016; Rafaluk-Mohr et al., 2018).

From this, we predict that variation in pathogen infection might limit the evolution of microbe-mediated protection. In the present study, we experimentally co-passaged *C. elegans* with protective *E. faecalis* and infected the host with evolutionary static pathogenic *S. aureus* at different intervals of host evolution. We also examined whether pathogen presence at the initial formation of the coevolving interaction is crucial to the evolution of protection. We show that enhanced microbe-mediated protection emerged out of novel coevolutionary host-microbe interactions and during pathogen infection, regardless of its temporal variability or the time point of first infection. Enhanced protection was only effective during pathogen infection. If hosts survived infection, they could recover and had the same longevity and reproductive output across treatments. These results thus suggest that even occasional pathogen infection can select for defensive mutualism, revealing the potential for this phenomenon to be widespread in nature.

## Materials and Methods

### Worm host and bacteria system

As a bacteriovore, *Caenorhabditis elegans* interacts constantly with a variety of bacteria either by feeding or hosting them (Cabreiro and Gems, 2013; Garsin et al., 2001; Schulenburg and Ewbank, 2004). Consequently, *C. elegans* is an established model for studying innate immunity (Gravato-Nobre and Hodgkin, 2005), as it can be infected with its natural (Jansson, 1994; Schulenburg and Ewbank, 2004) as well as opportunistic pathogens (Garsin et al., 2001; Tan et al., 1999). Most pathogens are taken up orally by the worm (Marsh and May, 2012), and some can proliferate and colonize the worm gut (King et al., 2016; Rafaluk-Mohr et al., 2018).

Naturally, *C. elegans* is a self-fertilising hermaphrodite (Brenner, 1974), but in this experiment obligate outcrossing worm populations (line EEVD00) with males and females (hermaphrodites that carry the *fog-2(q71)* mutation) were used (Theologidis et al., 2014). This lineage was generated by Henrique Teotonio (ENS Paris) and encompasses the genetic diversity of 16 natural worm isolates (Theologidis et al., 2014). Worms were kept on Nematode Growth Medium (NGM), inoculated with *Salmonella*, hereafter referred to as food. Worms were infected with the pathogenic *S. aureus* (MSSA476) (Holden et al., 2004), which is virulent and kills worm hosts by lysing the intestinal cells lining the gut wall (Sifri et al., 2003). Worms were exposed to *E. faecalis* (OG1RF) (Garsin et al., 2001), which was isolated from the human digestive system, but has been previously shown to colonize and proliferate in the host gut (Ford et al., 2017; King et al., 2016; Rafaluk-Mohr et al., 2018), where it provides protection.

### Experimental evolution - Design

Six single clones of *E. faecalis* (one for each of the six replicate populations) and a single population of *C. elegans* were the ancestors (hereafter referred to as the Ancestor) for all evolving populations. To account for potential differences in virulence, a stock of four clones of *S. aureus* was used for pathogen infections. Both *C. elegans* and colonising *E. faecalis* were allowed to evolve in presence of each other, while *S. aureus* was kept evolutionarily static. Infection with *S. aureus* was varied over host evolutionary time (indicated by purple in Table 1) to represent temporal heterogeneity in pathogen infection, including a range from always to every 2^nd^ generation, every 5^th^ generation, and never (Table 1). Moreover, we included differences in whether pathogens were present at the initial formation of the symbiotic interaction or later (2.1. vs. 2.2., and 5.1. vs. 5.2. in Table 1). Controls for lab adaptation were maintained for the host (No Protective-Microbe control, NPM in Table 1) and *E. faecalis* (No Host Control, NHC in Table 1).

**Table 1:**
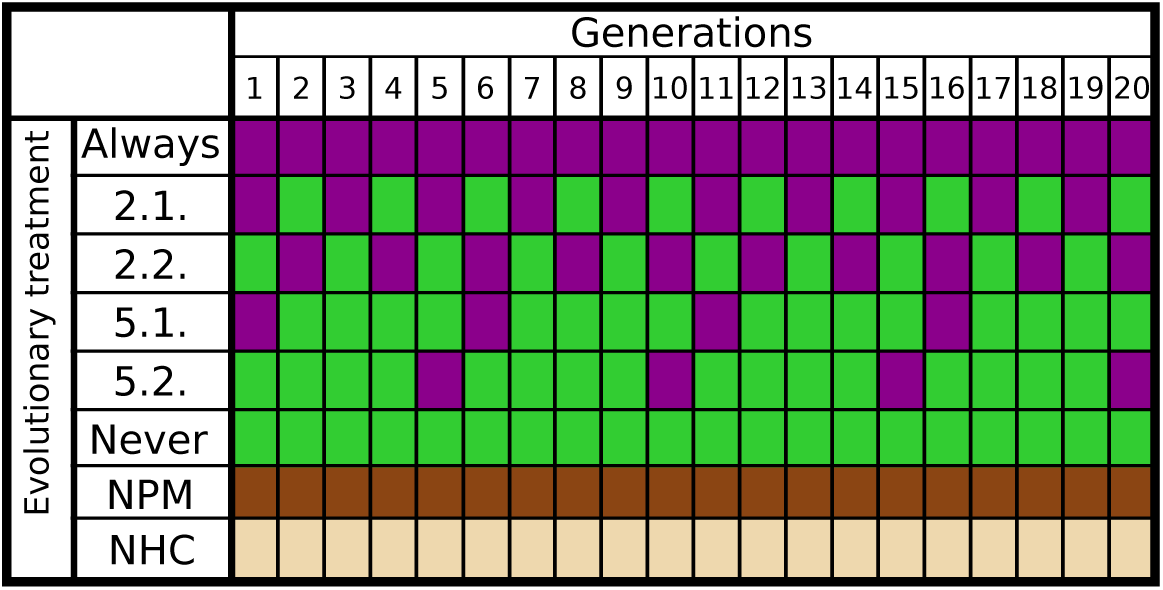
Experimental procedure for the evolution experiment. Columns indicate the number of experimental host generations (1-20), while rows show the eight treatments. Host generations were infected with *S. aureus* (purple) or given food (green), while constantly coevolving with *E. faecalis*. Two controls for lab effects on host evolution (dark brown, No Protective Microbe, NPM) and *E. faecalis* evolution (light brown, No Host Control, NHC) were also included, where the NPM treatment was only ever exposed to food alone. Each evolutionary treatment consisted of six independent evolutionary replicates.

### Experimental evolution - Culturing and passaging methods

At the start of each generation, worms were bleached as described previously and left in M9 buffer overnight for larvae to hatch (Stiernagle, 2006). Simultaneously, *E. faecalis* clones were cultured overnight in Todd-Hewitt Broth (THB) in 600µl at 30°C, while food was cultured overnight in LB broth. Subsequently, 9cm NGM plates were inoculated with 300µl of each overnight culture. Plates with freshly inoculated bacteria were dried at room temperature before approximately 1000 L1 worms were added to each NGM plate. After these plates dried at room temperature, they were transferred to a 20°C incubator and left for 48h. Simultaneously, a liquid culture of *S. aureus* was grown in THB from frozen stock, while a liquid culture of food was grown in LB, and both were incubated under shaking conditions at 30°C. The following day, 100µl of each overnight culture were spread on 9cm plates, *S. aureus* on Tryptone Soy Broth agar (TSB) plates and food on NGM plates and incubated at 30°C overnight. To transfer worms to the pathogen or food plates, nematodes were washed off the *E. faecalis* plates with M9 buffer and washed three times over small-pore filters to remove all externally attached bacteria, as previously described (Jansen et al., 2015; Papkou et al., 2019; Rafaluk-Mohr et al., 2018). Worms were infected with either *S. aureus* or exposed to food (Table 1) and left at 25°C for 24h. After this time, worms were then washed off the plates with M9 buffer once more to plate them on NGM plates seeded with food for laying eggs. Roughly 10% of these worms were crushed and plated on *E. faecalis* selective medium (TSB + 100mg/ml Rifampicin). The remaining worms were left on food plates for 48h to allow for egg laying.

To passage *E. faecalis*, roughly 100 *E. faecalis* colonies were picked and grown up shaking overnight in 600µl THB at 30°C, while worms were bleached and left to hatch overnight. This cycle was repeated for 20 experimental host generations.

All passaged worms and *E. faecalis* samples were cryopreserved at −80 °C. A proportion of the offspring of surviving worms were frozen in 40% DMSO, and 100µl of *E. faecalis* liquid culture was mixed with 100µl of glycerol before cryopreservation.

### Host survival and fecundity assays

All assays were conducted at the end of the evolution experiment on archived samples. Plates were randomized and fully encoded during each experiment to ensure the experimenter was blind to different treatments whilst collecting data.

Basic procedures were adopted from the experimental evolution, but with the following alterations to keep the assays feasible with higher accuracy when scoring dead and alive worms: 400 L1 worms were exposed to 200µl of food and *E. faecalis* on 6cm NGM plates, while 60µl *S. aureus* overnight culture was used to inoculate 6cm TSB plates.

To assess microbe-mediated protection of different combinations of worms and *E. faecalis*, 400 L1s were exposed to 50:50 mixtures of *E. faecalis* and food for 48h. Worms were then washed off these plates as described above and infected with *S. aureus* for 24h at 25°C. Survival in form of counting dead and alive worms was then scored.

To assess any long-term fitness consequences after protective microbe exposure and pathogen infection, long-term survival and fecundity were measured. Worms were exposed as described for the survival assays. Subsequently, five females and five males were picked onto 3cm food seeded NGM plates at 25°C and then transferred to new plates every 36h to avoid any confusion between offspring produced and original adults. At each time point, survival was scored. To measure fecundity, the number of worm eggs on the plates at 120h since bleaching were counted.

### Statistical Analysis

Statistical analyses were carried out with RStudio (Version 1.1.463 for Mac), graphs created with the ggplot2 package (Version 2.1.0) and edited with Inkscape (Version 0.91). All host survival and fecundity data were analysed with nested binomial mixed effects models (R package lme4), followed by a Tukey multiple-comparison tests (R package multcomp). Life-span data were analysed with Kaplan Meier Log Rank test with FDR correction for multiple testing.

## Results

Before the start of the evolution experiment, the starting conditions were tested. Confirming previous results, *E. faecalis* showed some spontaneous host-protective potential against *S. aureus*. Worms raised on *E. faecalis* and food survived better than those raised on food alone, independent of food or pathogen present at the later stage (General Linear Model, X^2^=10.205, df=1, p=0.001; Figure 1). Worms infected with *S. aureus* in later life survived worse than those being exposed to food (General Linear Model, X^2^=119.643, df=1, p<0.001; Figure 1). These results demonstrate the beneficial and protective effects for the host after exposure to the protective microbe *E. faecalis*.

**Figure 1:**
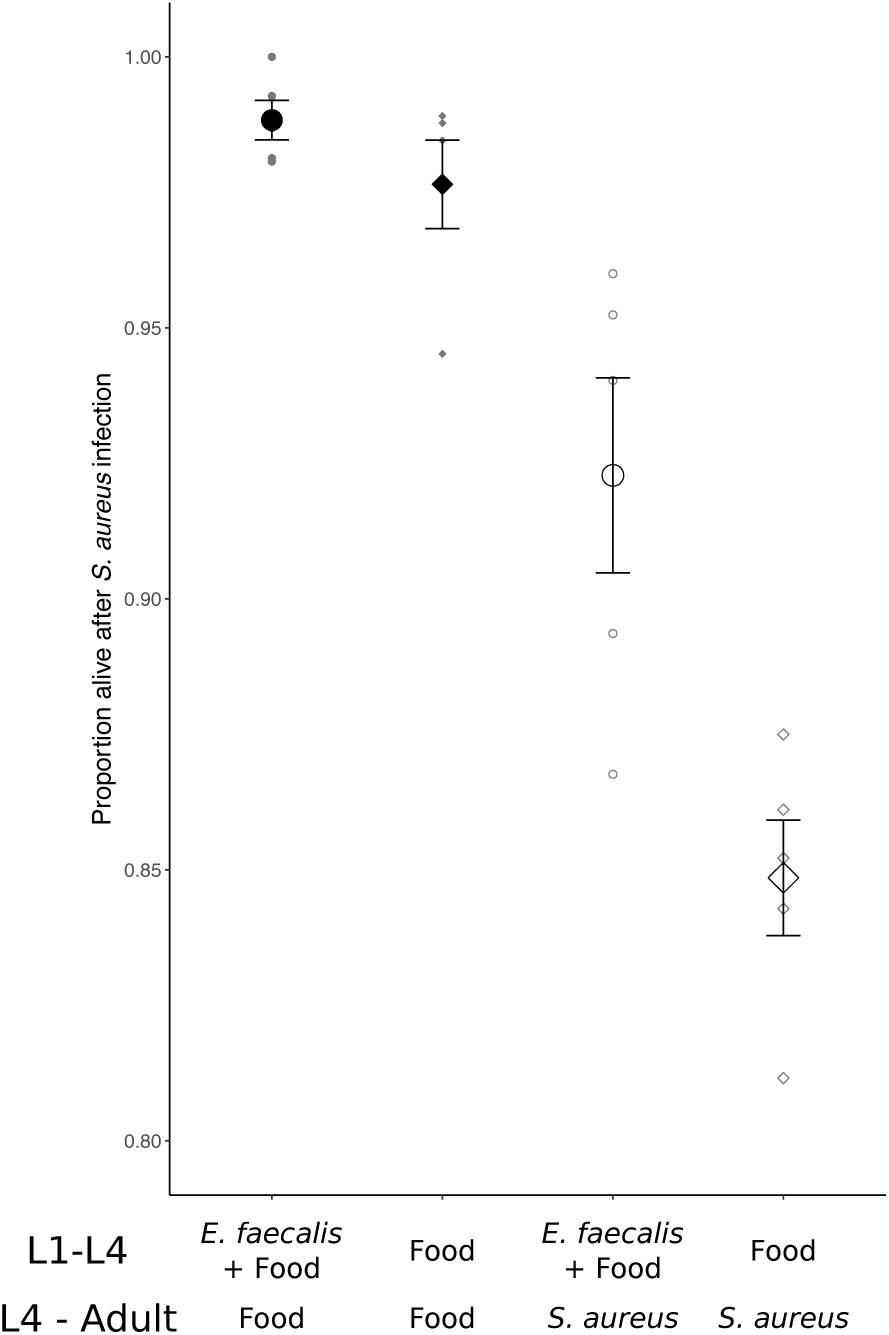
Host survival showing protective effects of *E. faecalis*. Early exposure of worms to *E. faecalis* (both ancestors) provides some degree of protection from the infection of *S. aureus*. 24h host survival levels reveal a benefit to *E. faecalis* colonisation independent of pathogen presence or absence. Circles indicate those treatment being exposed to *E. faecalis* and food in the earlier stage (L1-L4), while squares indicate food alone treatment in the earlier stage (L1-L4). Filled symbols indicate those treatments being exposed to food in the later stage, while open symbols indicate those treatments being exposed to the pathogen *S. aureus* in the later stage. Each symbol indicates the mean ± S.E of five replicates. Axis scales were chosen to be the same across all plots.

Infection with *S. aureus* over evolutionary time in the experiment led to the substantial enhancement of microbe-mediated protection, with the evolutionary background of the sympatric pair of host and *E. faecalis* having a significant impact on host survival (Mixed Effects Model, X^2^=42.479, df=4, p<0.001; Figure 2A). Higher microbe-mediated protection in comparison to the Ancestor occurred in all evolutionary histories involving pathogen presence across the temporal heterogeneity treatments in our evolution experiment (Always, 2.1. and 5.1.). However, this did not occur in the pathogen absence (Never) treatment. Host evolutionary history alone had a significant effect on host survival (Mixed Effects Model, X^2^=35.779, df=5, p<0.001; Figure 2B), but did not reveal the same pattern as for sympatric pairs. No effect of bacteria evolutionary history alone on infected host survival was observed (Mixed Effects Model, X^2^=3.2511, df=5, p=0.6613; Figure 2C). Taken together, enhanced microbe-mediated protection evolved only as a product of coevolution and pathogen presence for sympatric pairs; this occurred regardless of the temporal heterogeneity.

**Figure 2:**
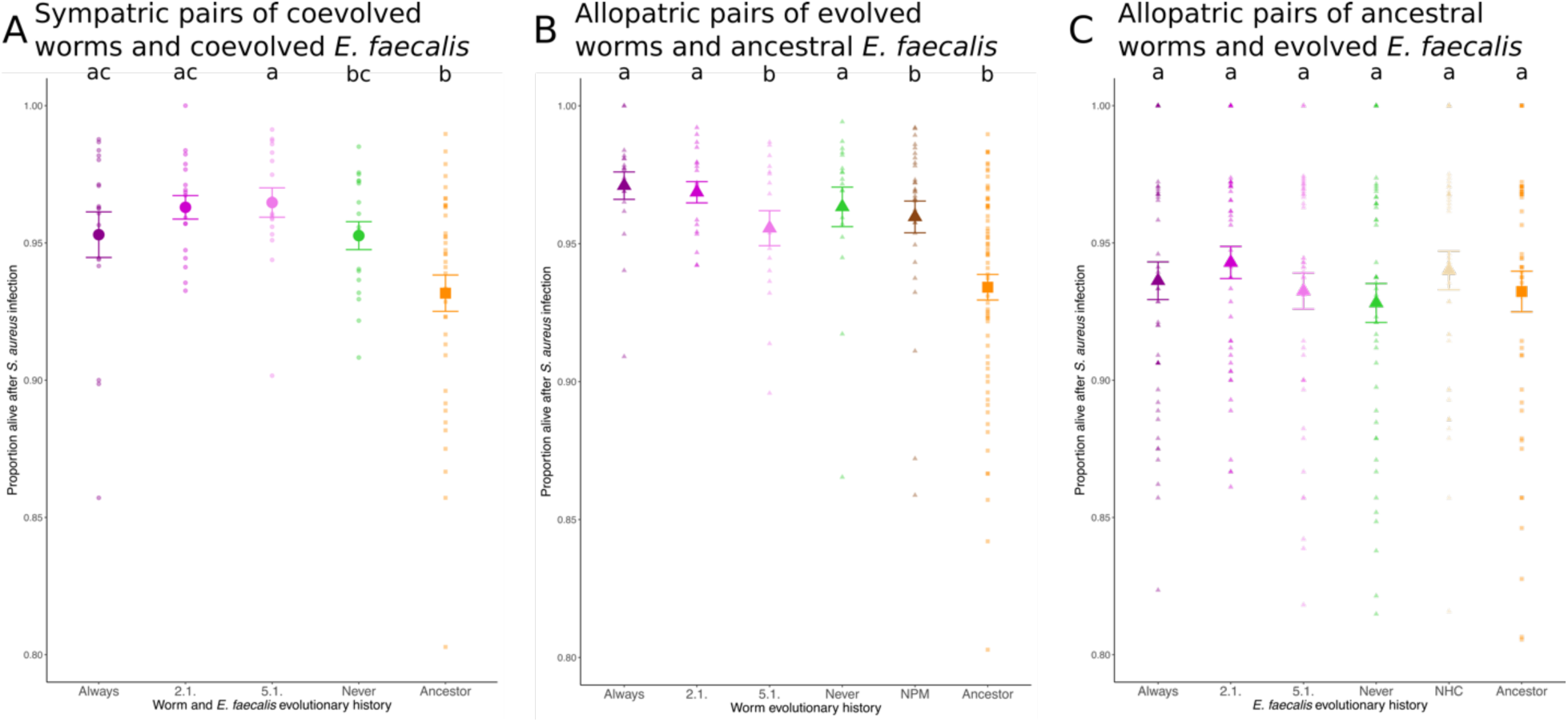
Host survival for coevolving sympatric and allopatric pairs of worms and *E. faecalis*. Microbe-mediated protection was assessed for (A) sympatric pairs of coevolved worms and *E. faecalis*, (B) allopatric pairs of evolved worms and ancestral *E. faecalis*, and (C) allopatric pairs of ancestral worms and evolved *E. faecalis*. Bigger symbols represent mean ± S.E. and consists of six biological replicates and four technical replicates. Smaller symbols indicate the data distribution. Circles indicate sympatric pairs of coevolved *E. faecalis* and worms, squares indicate ancestral pairs of *E. faecalis* and worms and triangles indicate allopatric pairs of *E. faecalis* and worms. Letters indicate results of a GLMM, followed by a Tukey Post-hoc Test. The same letter indicates no significant difference. Axis scales were chosen to be the same across all plots.

As an additional form of pathogen heterogeneity, the impact of the timing of initial pathogen infection on the evolution of microbe-mediated protection was investigated. An effect of different initial pathogen infection time points on host survival following pathogen infection was observed (Mixed Effects Model: X^2^=7.945, df=3, p=0.04716 Figure 3), although a Tukey Post-Hoc test revealed no significant differences (Table A1).

**Figure 3:**
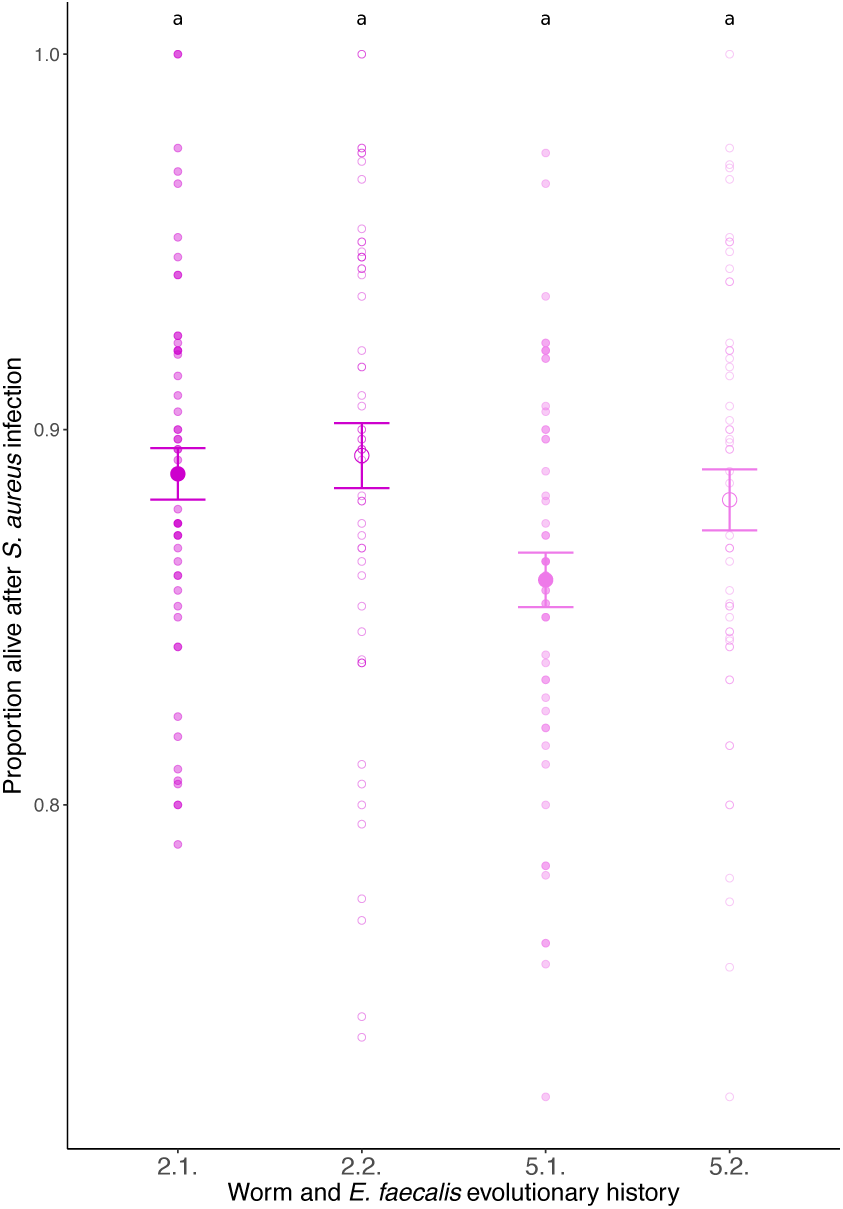
Host survival in evolutionary treatments differing in initial pathogen exposure time points. The time point of initial infection varied for infection to the pathogen every two generations (2.1. and 2.2) or every five generations (5.1. or 5.2.) but does not influence the outcome. Closed symbols indicate initial pathogen presence (Host generation 1), open symbols indicate later pathogen presence (Generation 2 for 2.1. and 2.2. and Generation 5 for 5.1. and 5.2.). Bigger symbols represent mean ± S.E and consists of six biological replicates and four technical replicates of the sympatric pairs. Smaller symbols indicate the data distribution. Letters indicate results of a GLMM, followed by a Tukey Post-hoc Test. The same letter indicates no significant difference. Axis scales were chosen to be the same across all plots.

Furthermore, we investigated the long-term consequences to hosts colonised by *E. faecalis* after 24h of pathogen infection. No significant differences were observed in the long-term survival post-infection of worm hosts colonised by their sympatric *E. faecalis* across treatments (Kaplan Meier Log Rank Test, FDR corrected, all comparisons p>0.05, Figure 4A). In addition, we did not find significant differences in fecundity among sympatric host-*E. faecalis* pairs (Mixed Effects Model, X^2^=3.9418, df=4, p=0.4278, Figure 4B).

**Figure 4:**
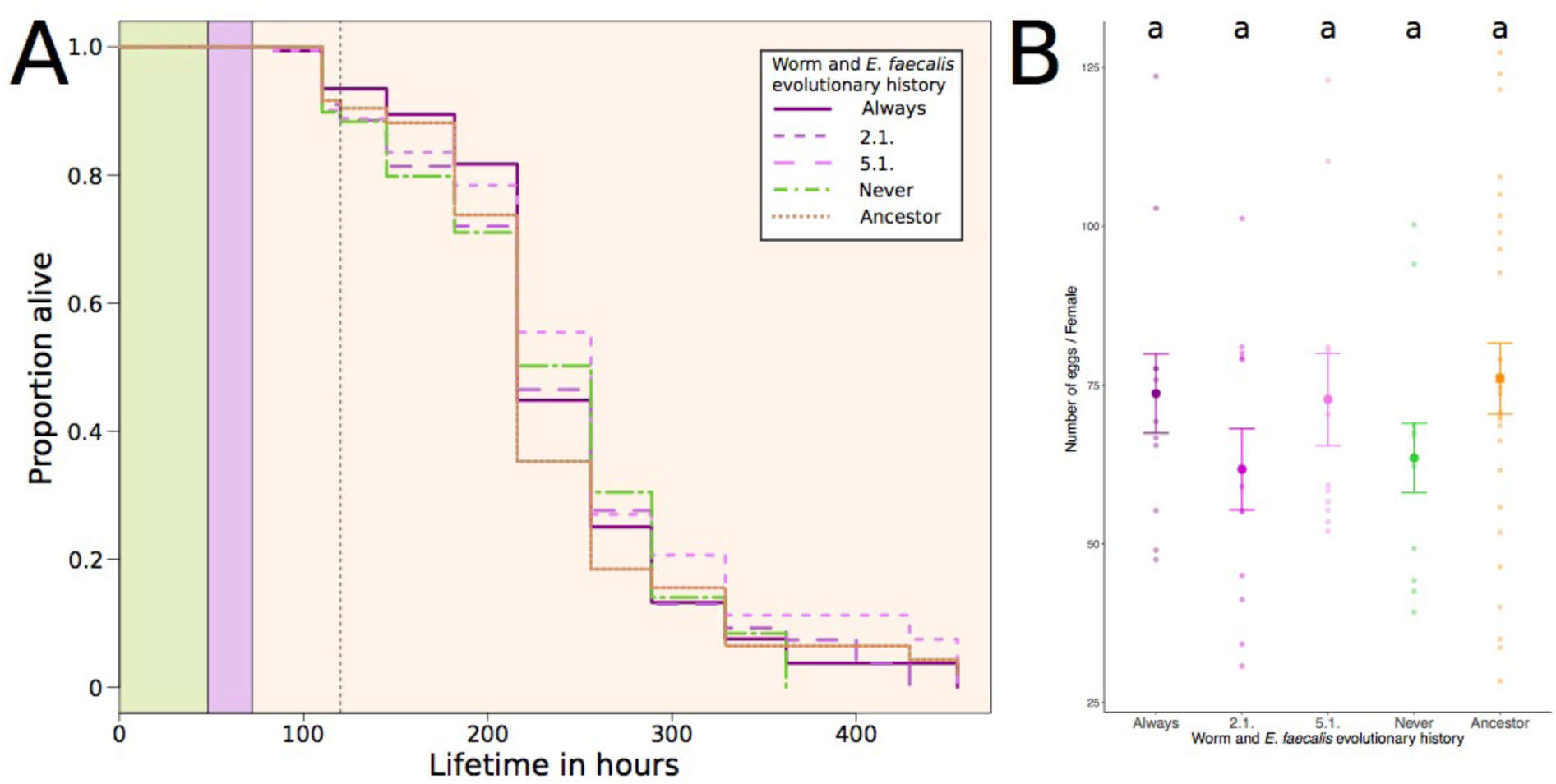
Long-term survival and fecundity of *E. faecalis-*colonised hosts that survived pathogen infection. (A) Long-term host survival was measured. Survival curves for sympatric pairs of worms and *E. faecalis* are shown as Kaplan-Meier estimates. Worms were exposed to *E. faecalis* and food (green), then to *S. aureus* (purple), and long-term survival was monitored on food (orange). The dotted line indicates the time point at which fecundity was measured. (B) Number of eggs/Female across sympatric pairs of coevolved worms and *E. faecalis*. Bigger symbols represent mean ± S.E. and consists of six biological replicates and four technical replicates. Smaller symbols indicate the data distribution. Circles indicate sympatric pairs of coevolved *E. faecalis* and worms, squares indicate ancestral pairs of *E. faecalis* and worms. Letters indicate results of a GLMM, followed by a Tukey Post-hoc Test. The same letter indicates no significant difference.

## Discussion

It has been shown that hosts receive the greatest benefits from protective microbes under constant pathogen infection. We hypothesized that variation in pathogen presence over time would limit the evolution of microbe-mediated protection due to the reduced benefits to the host and bacterial symbiont. In our study, enhanced pathogen defence emerged out of host-symbiont coevolutionary interactions only when pathogens were present, independent of the interval or initial presence of the pathogen. Notably, the ultimate strength of microbe-mediated protection that evolved was not impacted by the number of host generations between pathogen infections, the proportion of generations infected, or the presence of the pathogen at the first host-microbe interaction. These results suggest that resident microbes can be a form of transgenerational immunity against rare pathogen infections.

We found that microbe-mediated protection is maintained even in the prolonged absence of pathogen, but that pathogen presence is necessary for microbe-mediated protection to evolve, as previously hypothesized (Clay et al., 2005; King and Bonsall, 2017; Lively et al., 2005). This result is unlike previous work showing that the scale of heterogeneity in abiotic conditions can affect the strength of selection for traits in some symbiotic interactions (Harrison et al., 2013). This discrepancy is potentially due to costs in our symbiotic system being ameliorated (at least in terms of host survival) in well-provisioned hosts, as hosts are provided with food alongside *E. faecalis* and are thus rescued from starvation (also see (Dasgupta et al., 2019)). Although protective symbionts can incur costs (e.g., Vorburger & Gouskov, 2011) for their hosts, with potential for impacts on coevolutionary interactions (King and Bonsall, 2017), it is possible that potential costs of bacterial colonisation might be only detectable when hosts are stressed (Lively, 2006) or that the costs were not strong enough for us to detect(Little et al., 2002). Different measures of cost remain to be explored (e.g. lifespan in the complete absence of a protective microbe and a pathogen). Higher protection also does not always come with higher costs, as found in the black bean aphid-*Hamiltonella defensa* interaction (Cayetano et al., 2015). Thus, protective traits in an organism’s commensal microbiota could be selected for under pathogen infection and easily maintained in subsequent uninfected generations.

Microbe-mediated protection was strongest between sympatric pairs when pathogens were present over evolutionary time, consistent with previous findings (Rafaluk-Mohr et al., 2018). In our study, protection emerged during coevolution after only 20 host generations, and not due to the independent evolution of either interacting species, but due to the coevolution of both species (King and Bonsall, 2017). The time-scale of these interactions is short compared to the longer shared evolutionary histories shared by other defensive mutualisms (Jousselin et al., 2003; Quek et al., 2004; Shoemaker et al., 2002). Nevertheless, our findings reveal the potential for microbe-mediated protection to become enhanced during the formation of a coevolving host-microbiota relationship.

In conclusion, our results show that enhanced protection in host-microbe interactions can rapidly evolve and be maintained even under infrequent pathogen infection, suggesting that resident microbes can be a form of stable, transgenerational immunity. The protective benefit of an organism’s microbiota might remain undetected for several host generations until pathogens re-emerge. Future research on the failure of pathogens transmit within host populations should consider the contribution of the protective microbiota to prevent disease spread.

## Authorship Contributions

KCK and AK conceived the project. KCK, MBB, and AK designed the project. AK performed the evolution experiment and all subsequent assays. AK and MBB performed the statistical analysis. AK, KCK and MBB wrote the manuscript and agreed on the final version.

## Acknowledgments

We would like to thank the King group for help and support during the performance of the evolution experiment, especially Charlotte Rafaluk-Mohr and Maria Ordovas-Montanes. AK was supported by a fellowship from the “Studienstiftung des Deutschen Volkes”. KCK is grateful for a Leverhulme Trust project grant (RPG-2015-165) and ERC Starting Grant (COEVOPRO 802242).

## Conflict of Interest Statement

The authors declare no conflict of interest.

## Appendix

### Experimental Procedure

**Figure A1:**
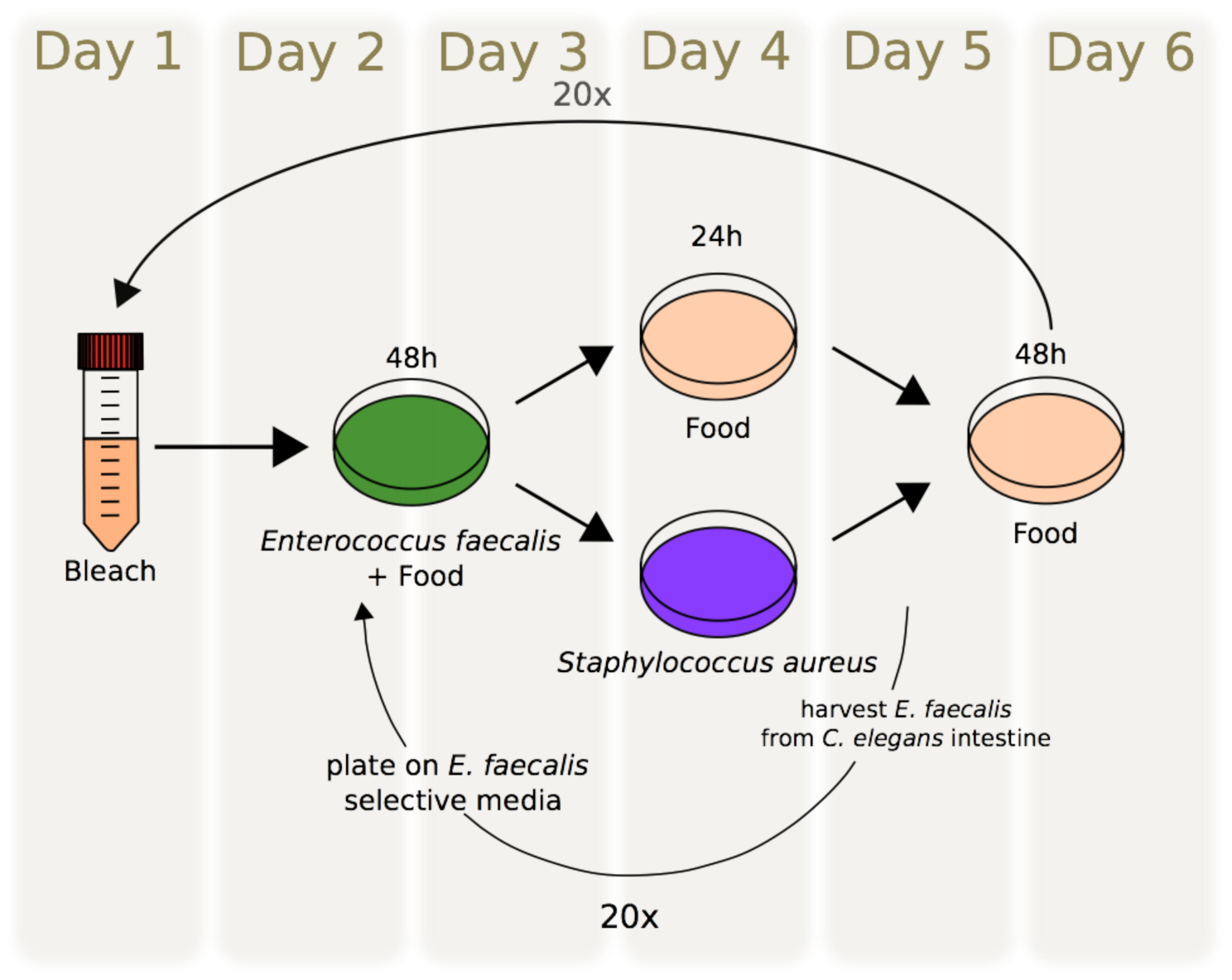
The experimental procedure of the evolution experiment in detail.

The procedure of the evolution experiment can be seen in Figure S1 and will be the following:

1. On the first day of each generation, worms were bleached(Stiernagle, 2006). During this step worms were removed from the plates by washing with M9 buffer. Afterwards a 50:50 mixture of 10% NaClO and 5M NaOH was used to bleach all the remaining bacteria in the solution and to release eggs from bleached adult worms. Only eggs survive this step and were then left in the M9 buffer over-night. All bleached eggs hatched overnight, but do not develop any further and thus lead to increased synchronisation for each worm population. Every bleach solution was plated out on a TSA plate to control for carried over contaminations. 100 colonies of *E. faecalis* were picked from the plates that were grown on *E. faecalis* selective medium for 48 hours at 30°C. These colonies were picked into 600μl of THB and then grown up over-night. Simultaneously, a single colony of *Salmonella* food was picked and added to 25ml of LB broth to be grown under shaking conditions over night.
2. On the second day of each generation, the overnight shaken and hatched worms were exposed to 600μl of a 50:50 mixture of *E. faecalis* and food. At this step the population size was adjusted to only contain 1000 individuals. Worms remained on these plates with *E. faecalis* for 48 hours.
3. On the third day of each generation, TSA plates to which worms were going to be exposed on day 4 were inoculated with 100μl of either *S. aureus* or food and were incubated at 30°C overnight.
4. On the fourth day of each generation, worms were washed off the plates seeded with *E. faecalis* by filter tip washing. For this purpose, worms were washed off the plates with twice 1.5ml of M9 buffer +1% Triton X100, as previously described(Jansen et al., 2015; Papkou et al., 2019; Rafaluk-Mohr et al., 2018).This worm and bacteria suspension was spun down for 1 minute, 1 ml of the supernatant was discarded and the rest of the pellet was pipetted on the top of a filter of a filter tip and spun down for 3 min. Worms were left on the top of the filter were washed with 400μl of M9 three times, before being re-suspended in 100μl of M9 to bring onto plate. During this method, most of the externally attached bacteria are washed off the worms to ensure that worm survival can be attributed to gut colonization of *E. faecalis* and not external attachment of the protective microbe. *Enterococcus faecalis* remains in the worm’s intestine and will establish a protective effect. After worms were transferred to the plates containing either *S. aureus* or food, all plates were moved to 25°C.
5. On the fifth day of each generation, worms were washed off the plates again by filter tip washing (as described for day 4 and previously (Jansen et al., 2015; Papkou et al., 2019; Rafaluk-Mohr et al., 2018)). Worms were left on plates seeded with food at 20°C for 48 hours to lay eggs. The amount of transferred bacteria (either *S. aureus* or food can be neglected to have any influence on the further development of the worms. These plates were then used for bleaching on day 1 of the following generation. 10% of the worm mixture was separated and used to isolate *E. faecalis*. For this purpose, the suspension of worms was crushed and then plated on TSA plates with Rifampicin, as the *E. faecalis* strain carries a Rifampicin resistance. The plated gut content was allowed to grow at 30°C for 48 hours.

#### Statistical results

**Table A1:**
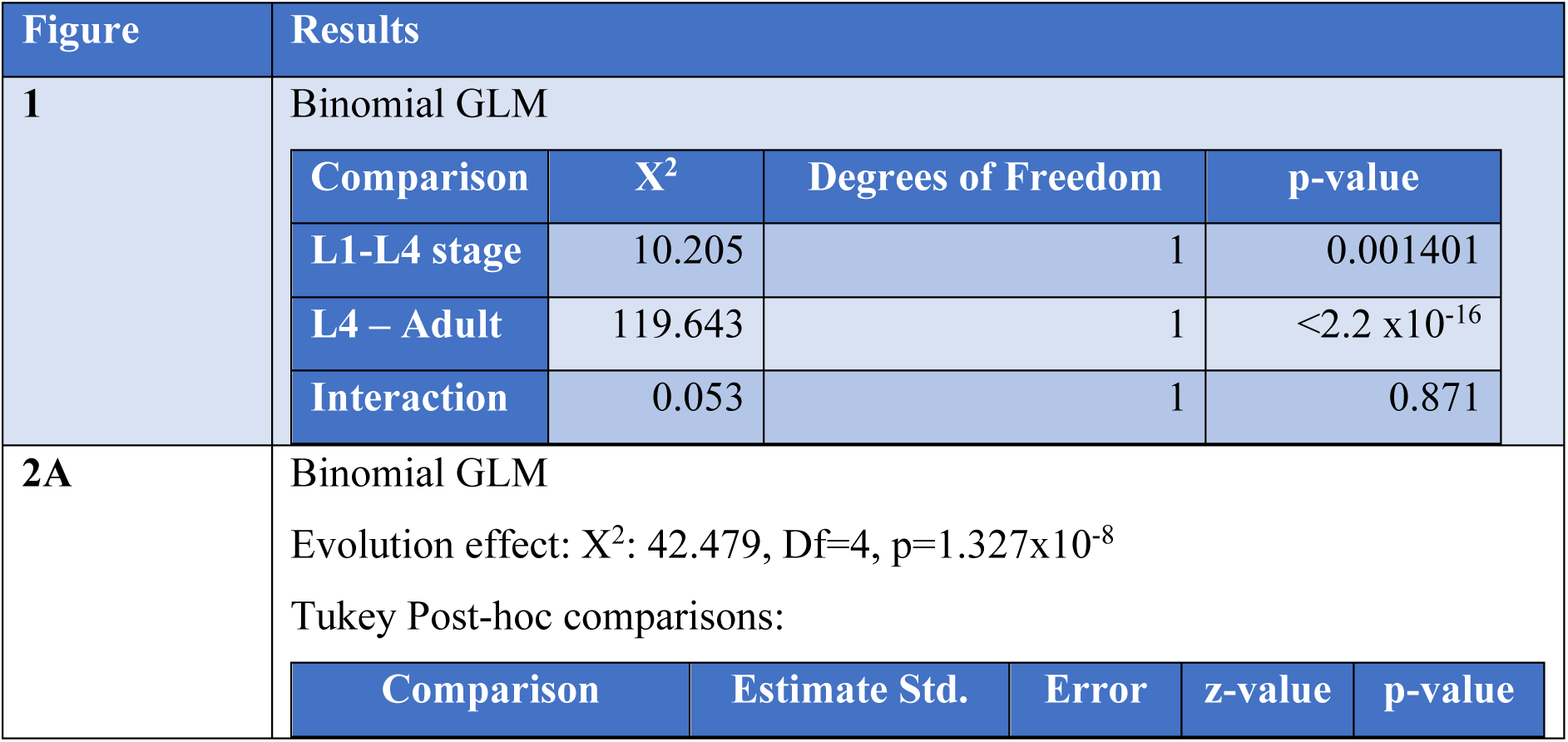
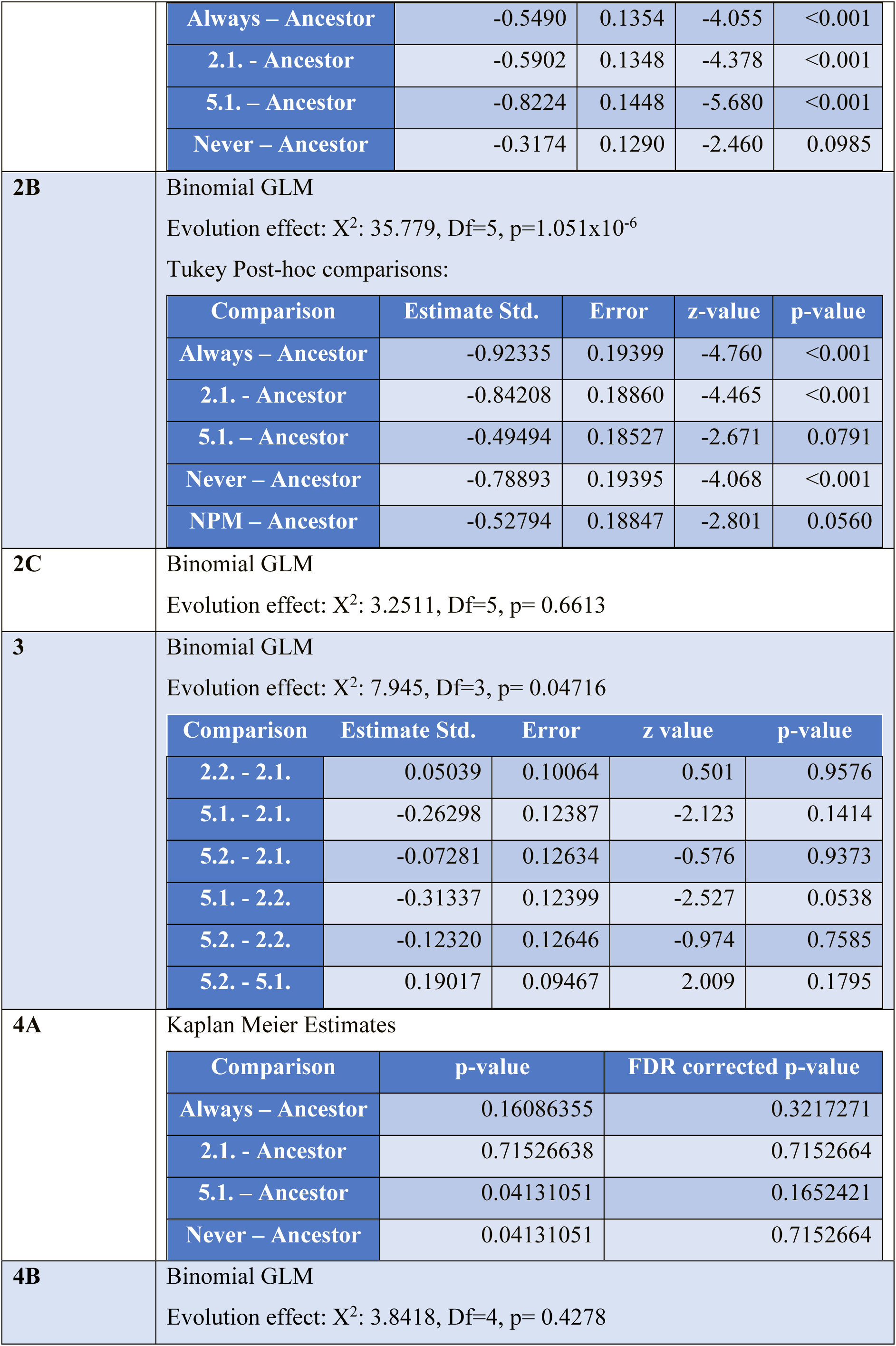
All statistical results summarized, including the statistical test, the specifics associated with each test, the relevant degrees of freedom and p-values.

## Notes

### Competing Interest Statement

The authors have declared no competing interest.

### Summary of Updates

We have clarified the wording and the wider application of our results. We have adjusted the graphs slightly, so results are easier visible. Further explanation for statistical differences and experimental procedure (particularly for the Evolution Experiment) have been added to the Supplemental Material

